# A wide foodomics approach coupled with metagenomics elucidates the environmental signature of Protected Geographical Indication potatoes

**DOI:** 10.1101/2022.10.12.511727

**Authors:** Anastasia Boutsika, Michail Michailidis, Maria Ganopoulou, Athanasios Dalakouras, Christina Skodra, Aliki Xanthopoulou, George Stamatakis, Martina Samiotaki, Georgia Tanou, Theodoros Moysiadis, Lefteris Angelis, Christos Bazakos, Athanassios Molassiotis, Irini Nianiou-Obeidat, Ifigeneia Mellidou, Ioannis Ganopoulos

## Abstract

The term “terroir” has been widely employed to link differential geographic phenotypes with sensorial signatures of agricultural food products, influenced by agricultural practices, soil type and climate. Nowadays, the Geographical Indications labeling has been developed to safeguard the quality of plant-derived food that is linked to a certain terroir and is generally considered as an indication of superior organoleptic properties and phytochemical profile. As the dynamics of agroecosystems are highly intricate, consisting of tangled networks of interactions between plants, microorganisms, and the surrounding environment, the recognition of the key molecular components of terroir fingerprinting remains a great challenge to protect both the origin and the safety of food commodities. Furthermore, the contribution of microbiome as a potential driver of the terroir signature has been underestimated until recently. Herein, we present a first comprehensive view of the multi-omic landscape related to transcriptome, proteome, epigenome, and metagenome of the popular Protected Geographical Indication potatoes of Naxos.

## Introduction

Nowadays, given the globalization as well as the numerous technological developments and innovations that govern the food market, consumer’s expectations have been tremendously increased in terms of information reliability. Therefore, the food industry and governments should be provided with valid analytical methods and regulatory frameworks to ensure food certification via the reliability of food labels taking into account consumer’s requirements. In the past decades, a plethora of deception incidents have been occurred, bewildering the food market (Braconi et al., 2021). Consequently, international operations have developed a profound interest in preventing food fraud and securing food authentication. One of the most common frauds involves selling low-quality food products at high prices. Accordingly, food labels could include fabricated geographical origin or genetic identity, as well as inaccuracies in the production process (Braconi et al., 2021). Although food fraud appears to be price-related, in some cases ingredients dangerous to human health and allergens may be included in these food products, endangering consumer safety (Braconi et al., 2021).

To prevent fraud of food products, scientists have developed several rapid, reliable, and efficient -omics technologies for certification and identification, including genomics, epigenomics, transcriptomics, proteomics and metabolomics. The term ‘Foodomics’, concerning the study of Food and Nutrition in combination with -omics technologies, was initially launched in 2009 (Capozzi and Bordoni, 2012). Despite the short time that the ‘Foodomics’ domain exists in the scientific community, a wealth of technologies has been developed aiming to study the quality, the origin, and the safety of human nutrition (Ahmed et al., 2022). The most popular and intriguing challenge in food research is the validity of food labels, especially on products designated as Protected Designation of Origin (PDO) or Protected Geographical Indication (PGI), according to the EU geographical indications system for food quality.

Potato (*Solanum tuberosum* L.) was a fundamental species in human nutrition especially in the context of an ever-increasing population. Nowadays, potato remains one of the most popular and crucial nongrain food crops, occupying a prominent place in the agenda of global food security (Pearsall, 2008; Spooner et al., 2005). According to Plant Production and Protection Division, 2009, it is estimated that two billion people worldwide are closely associated with potato cultivation for nutritional or income reasons, rendering it as “Food for the Future” (Ortiz and Mares, 2017).

Using potato as a plant model, in this proof-of-concept study we present the first multi-omics analysis across genome-wide DNA methylation, RNA sequencing and quantitative proteomics to obtain the molecular portrait of the famous PGI potatoes of the Naxos Island at harvest and after storage. We also employed a metagenomic approach to discriminate potato tubers produced in diverse regions based on the distinct microbiological patterns, which in turn were coupled with the -omics datasets. Through this approach, key environmental-derived molecular factors through the dynamics of causal models were revealed.

## Results

### Bacterial community dynamics in periderm of tubers

The tubers are harvested and traded with soil residues in the periderm, which makes them ideal for using the microbial community profiling as potential signatures in PGI certification. Thus, to detect distinct differences in the tuber bacterial profiles cultivated in the two different agroecosystems, bacterial 16S rRNA gene amplicon sequencing was performed in the tubers obtained at harvest and post-harvest (storage) (Figure 1, Table S1). The alpha-diversity highlighted greater species richness in the tubers from Naxos. Regarding species diversity, the microbial differences were only evident at post-harvest period, with tubers from Naxos exerting higher microbiome diversity (Figure 1B). These results may be indicative of the more rich and diverse microbial community of the PGI potato, especially after storage. By contrast, tubers from Lakoma (herein served as control), seemed to be dominated by fewer microbial species. Overall, microbiome communities at harvest of the Naxos tubers were dominated by *Lysobacter, Neobacillus* and *Priestia*, whilst the most abundant microbiota in the Lakoma tubers were *Rhizobium, Devosia, Sphighomonas* and *Rhodoligotrophos* (Figure 1C). Similar taxa were recorded as abundant for tubers at post-harvest, with several of them being widely recognized as plant growth-promoting rhizobacteria. Our data also demonstrated that, regardless the collection site, tubers at post-harvest maintained their microbial community profiles at the genus level to a great extent, providing a tool for PGI signature.

**Figure 1.**
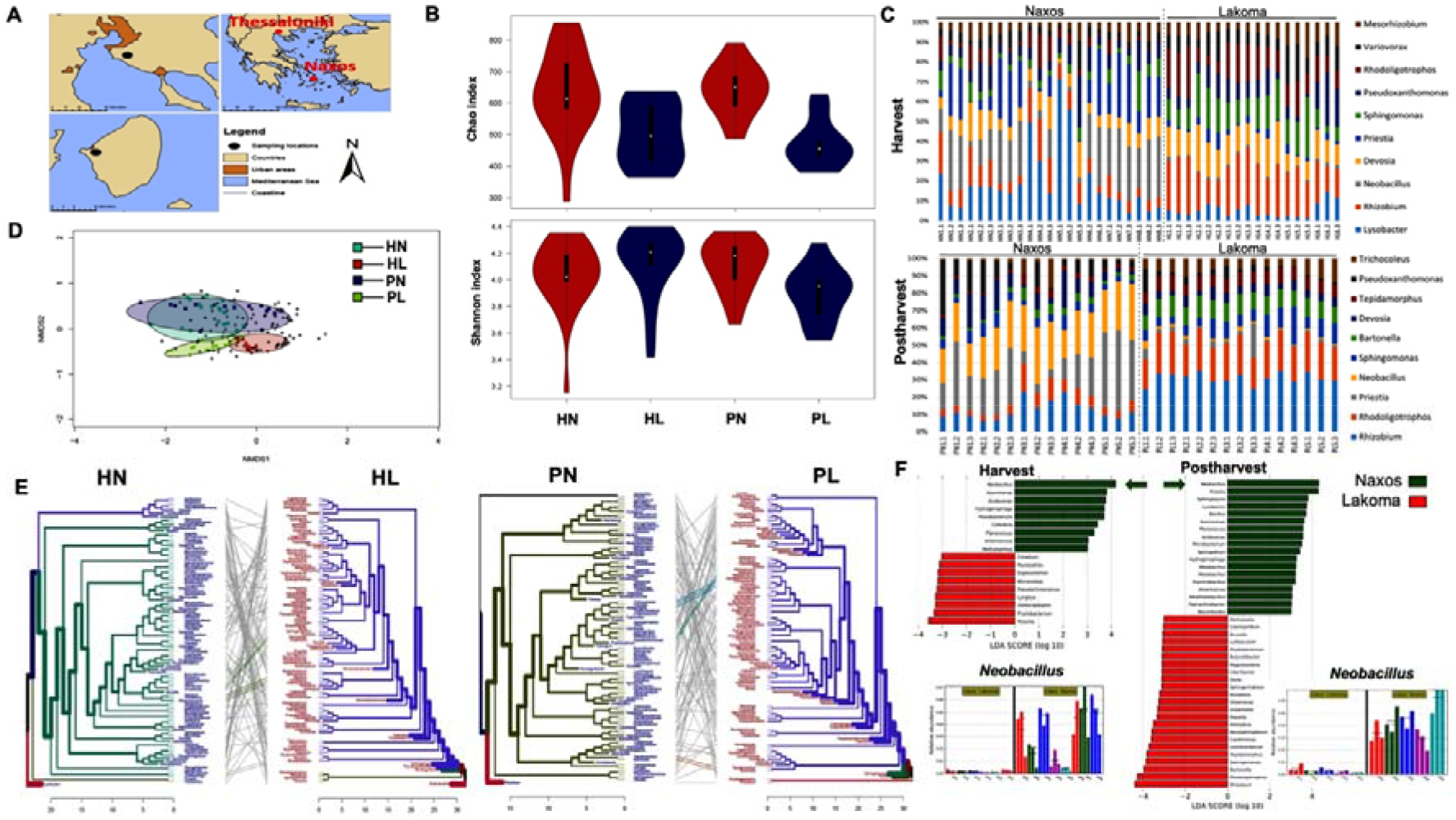
(A) Sampling locations of tubers in Naxos (PGI potatoes) and Lakoma (control potatoes). (B) Box plots of alpha-diversity (Chao and Shannon indices) of microbiome residing in the tubers from the two regions, at harvest and post-harvest. (C) Distribution of the top 10 most abundant taxa of tubers microbiota at the level of genus. (D) Microbiome profiles in the tubers obtained from the two regions analyzed by NMDS using the Bray–Curtis distance matrix. (E) Tanglegrams showing concordance between bacterial dendrograms based on community similarities (Bray–Curtis distance) derived from 16S rRNA gene sequences from tubers of the two regions. (F) Histogram of LDA value distribution of taxa at the genus level with significant differences in abundance between groups N: Naxos; L: Lakoma; H: Harvest; P: Post-harvest. Data obtained from Table S1.

The NMDS analysis with the Bray–Curtis dissimilarity (β-diversity) indicated that tubers from the two different agroecosystems were grouped separately from each other (Figure 1D), validating our hypothesis that the collection sites have distinct microbial community composition. In addition, tanglegrams between bacterial dendrograms showed that the bacterial structures between the two agroecosystems were dissimilar both at harvest and post-harvest (Figure 1E). The LEfSe analysis detected 18 and 41 bacterial clades in the tubers at harvest and post-harvest, respectively, discriminating the terroir-specific microbial communities in the two geographic regions. The dominant bacteria genus at harvest were *Neobacillus* and *Massilia* in Naxos and Lakoma, respectively. At post-harvest, candidate biomarkers belong to the genus of *Neobacillus* and *Priestia* for tubers of Naxos, or of *Rhizobium* and *Rhodoligotrophos* for tubers of Lakoma. These potential biomarkers were associated with each terroir, revealing their geographic-origin dissimilarities. Interestingly, one genus identified as bacterial biomarker of Naxos, *Neobacillus*, was detected both at harvest and at post-harvest. Therefore, this genus not only seemed to be abundant in the PGI potatoes, but it also remained abundant after storage, dominating the microbial community of potato tubers, and thus representing an excellent putative ‘terroir’ biomarker for traceability.

### Individually constant methylated genes, expressed transcripts, and level of proteins in PGI potato tubers

To gain a comparative insight into how different environments built PGI signature, large scale -omics technologies, i.e., epigenomics, transcriptomics, and proteomics, were applied.

#### Epigenetic marks of the potatoes from the two geographic regions

Plant epigenetic profile can be highly dynamic and plastic in diverse environmental conditions (Dalakouras and Vlachostergios, 2021), therefore we compared the DNA methylome of the tested potatoes (PGI and control) at harvest and at post-harvest using whole genome bisulfite sequencing (WGBS). Chromosome level analysis of differentially methylated regions (DMRs) at harvest and post-harvest revealed hypomethylation and hypermethylation events in both gene regions and transposable elements (TEs) (Figure 2A). DMR methylation level cluster heatmap and violin plot highlighted even further these differences (Figures 2B and 2C). Focusing on DMR-associated genes (DMGs) exhibiting hypermethylation or hypomethylation, we could detect at least 13 and 29 DMGs at harvest and post-harvest, respectively (Figure 2D). When analyzing the distribution of DNA methylation among various gene features, most DNA methylation (especially at CG context) was recorded in the promoter and intron sequences rather than in exon and UTR sequences (Figure 2E). The distribution of DNA methylation in upstream 2K and downstream 2K regions, we observed that while gene bodies exhibited high mCG/CG but low mCHH/CHH ratio, the opposite took place in upstream 2K and downstream 2K sequences (Figure 2F). Based on Kyoto Encyclopedia of Genes and Genomes (KEGG) pathway enrichment analysis, DMGs were mainly assigned to metabolic pathways and biosynthesis of secondary metabolites (Figure 2G). A heatmap of DMGs displaying simultaneously all three sequence contexts (CG, CHG, CHH) allowed a broader overview of the epigenetic plasticity events at both examined periods between the two regions (Figure 2H). For instance, the hypermethylation of a putative DETOXIFICATION 18 (Soltu.Atl.10_4G001390) has been detected at harvest, whereas hypomethylation of three chloroplastic plastoglobulins-1 (Soltu.Atl.08_1G001340, Soltu.Atl.08_3G001840, Soltu.Atl.08_3G001850) has been determined at post-harvest.

**Figure 2.**
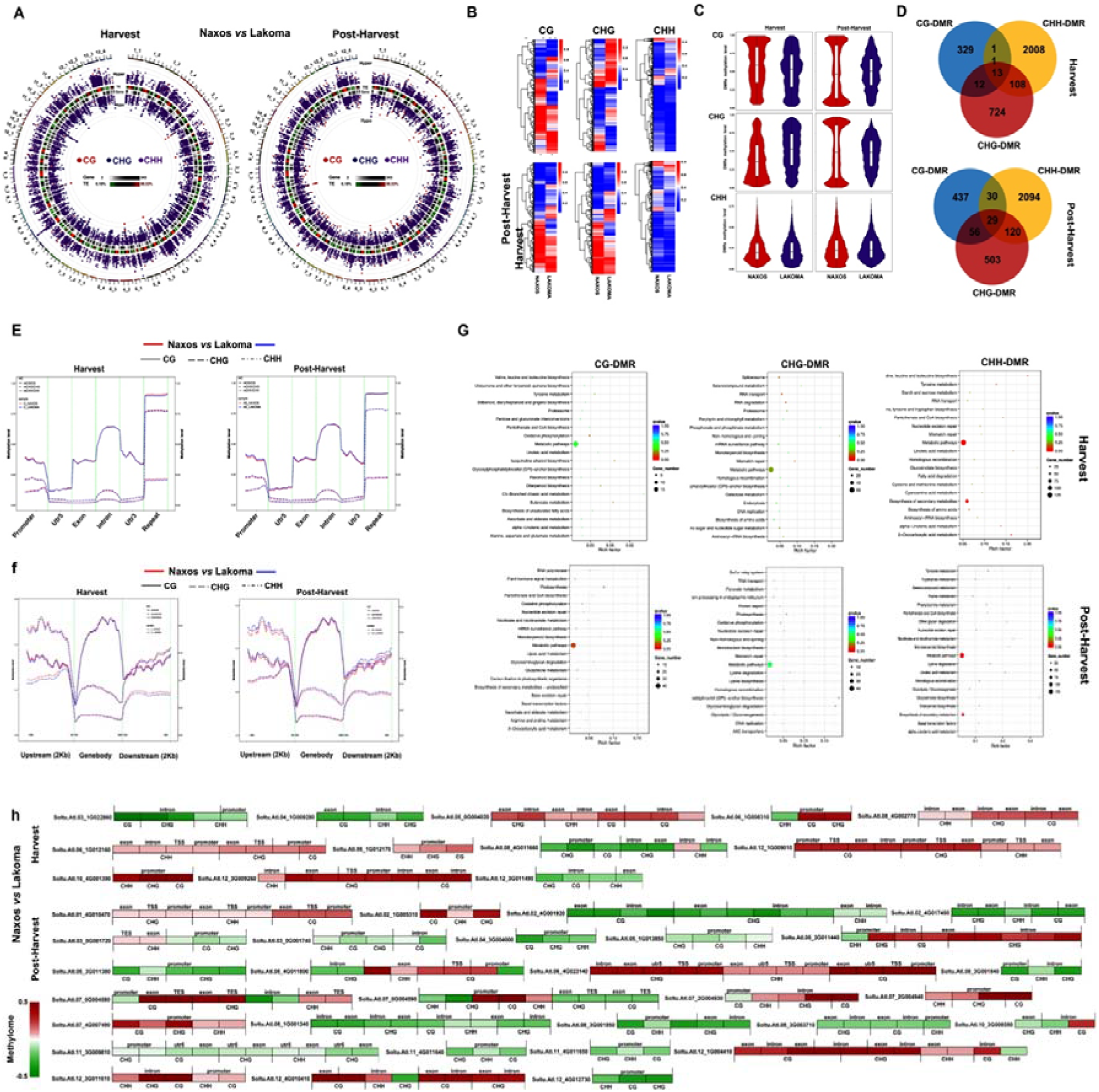
Differential methylation region (DMR) of the tubers harvested from Naxos (PGI potatoes) and Lakoma (control potatoes), at harvest and at post-harvest. **(A)** Circos plot for DMR condition in three contexts (CG, CHG, CHH). The circos plot represents (from outside to inside): (i) Hyper DMR statistical value: log5 (areaStat); the higher and bigger the point, the larger differences between two groups. (ii) TE, the heatmap of percentage of repeat element. (iii) Heatmap of gene density. (iv) Hypo DMR statistical value: log5 (areaStat); the higher and bigger the point, the larger differences between two groups. **(B)** Cluster heatmap for DMR methylation level in three contexts (CG, CHG, CHH). The x-axis is the comparison group name, the y-axis is the methylation level and cluster results. **(C)** Violin plot for DMR methylation level in three contexts (CG, CHG, CHH). The x-axis is the comparison group name, the y-axis is the methylation level. **(D)** Venn plot of DMGs in three contexts (CG, CHG, CHH). (**E**) Methylation level distribution at functional genetic elements in three contexts (CG, CHG, CHH). The x-axis is the functional genetic elements the y-axis is the methylation level. Left label is methylation level in non-CG context; the right label is methylation level in CG context. **(F)** Methylation level distribution at up/downstream 2kb and gene body in all three contexts (CG, CHG, CHH). The x-axis is the functional genetic elements, the y-axis is the methylation level. Left label is methylation level in non-CG context, the right label is methylation level in CG context. **(G)** KEGG enrichment scatter plot for DMR genes in all three contexts (CG, CHG, CHH). The x-axis represents Rich factor, and the y-axis represents pathway name. The size of points stand for DMR-related genes counts and the colors stand for different q-values range. **(H)** Heatmap of DMGs genes in all three contexts (CG, CHG, CHH). Red indicates hypermethylation and green hypomethylation in Naxos. Data obtained from Table S2.

#### Transcriptomic profiles of the potatoes from two distinct geographic regions

RNA-seq experiment was also conducted for the same samples as used for methylation analysis (Table S3). TPM-normalized values for each transcript were hierarchical-clustered and used to generate a heatmap that clearly shows a distinct expression pattern acceding to region and especially to stage (Figure 3A). Venn diagram showed genes that were commonly and exclusively modulated by the different environments and stages in potato (Figure 3B). For example, 940 and 947 genes were differentially expressed in ‘Naxos’ potatoes compared to the ‘Lakoma’ ones at harvest and at post-harvest, respectively (Figure 3B). Interestingly, we found 31 commonly expressed DEGs between Naxos and Lakoma, at harvest and at post-harvest, including cysteine protease inhibitor (Soltu.Atl.03_3G022650) and Kunitz trypsin inhibitors (Soltu.Atl.03_4G014820).

**Figure 3.**
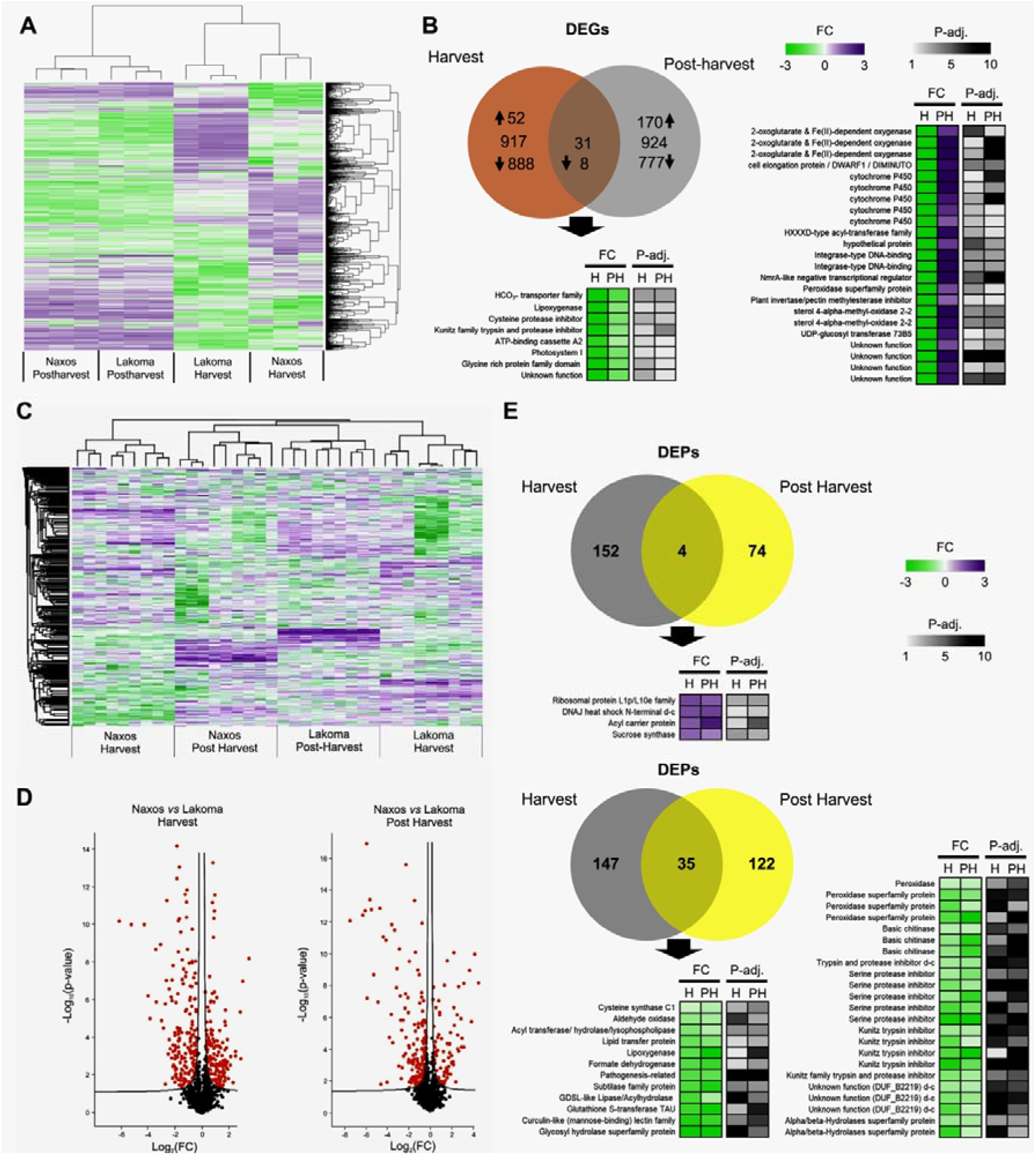
Transcriptome and proteome profiles of tubers in the regions of Naxos (PGI potatoes) and Lakoma (control potatoes). (A) Hierarchical cluster analysis of transcriptomic data in tubers of two regions at harvest and at post-harvest. (B) Venn diagrams of differentially expressed genes (DEGs) between Naxos and Lakoma at harvest (H) and post-harvest (PH). For the common DEGs between Naxos and Lakoma at the two stages, heatmaps representing the fold change (FC) in FPKM values of the up- and down-regulated genes at H and at PH are also provided. (C) Hierarchical cluster analysis of proteomic data in tubers of the two regions at H and at PH. (D) Volcano plots of Lakoma *vs* Naxos at H and PH. (E) Venn diagrams of differentially expressed proteins (DEPs) between Naxos and Lakoma at H and at PH. Heatmaps of the commonly increasing or decreasing DEPs in Naxos *vs* Lakoma are also provided. Data obtained from Table S4-S5.

#### Protein signature of the PGI potato tubers

To interpret the proteomic data in a PGI context, we focused only on proteins that constantly accumulated in the tubers of Naxos (PGI) at both stages. Proteomic data were clustered via Hierarchical Cluster Analysis demonstrating a distinct separation mainly between regions and secondly between stages (Figure 3C). Volcano plots revealed 156 and 78 differentially expressed proteins (DEPs) that were increased in Naxos *vs* Lakoma, at harvest and post-harvest, respectively (Figures 3D, 3E). Similarly, 182 and 157 DEPs decreased in Naxos *vs* Lakoma, at harvest and post-harvest, respectively (Figures 3D, 3E). Four proteins were increased in Naxos compared to Lakoma in both stages, being annotated as ribosomal protein L1p/L10e family (Soltu.Atl.11_1G013990.1), DNAJ heat shock N-terminal d-c (Soltu.Atl.03_1G024300.1), acyl carrier protein (Soltu.Atl.06_2G011230.1) and sucrose synthase (Soltu.Atl.09_1G015370.3). Moreover, 35 proteins (i.e., serine protease inhibitors, peroxidases, basic chitinases and Kunitz trypsin inhibitors) were decreased in both stages of Naxos compared to Lakoma (Figure 3E).

### Transcriptome-based pairwise co-expression analysis across multi-omics datasets reveals molecular hallmarks in PGI potatoes

There is a large interest in networked food science experiments for characterizing PGI signatures at molecular scale. Consequently, our work presents a pipeline system for pairwise integration and transcriptome-based co-expression analysis of epigenomic, transcriptomic, and proteomic data (Figure 4, Tables S6-S9). Our findings indicated that Pearson correlation coefficients showed negative values between the transcriptome and methylome datasets for both promoter (53.3%) and gene (50.58%), with 1% and 1.16% significant values (Figures 4A, 4B) while between the transcriptome and proteome datasets, positive correlation values were mostly observed (52.67%), with a 4.94% out of them being significant (Figure 4C). Methylation datasets of Naxos and Lakoma were positively correlated only at harvest stage (Figures 4D, 4E), while their proteomes exhibited positive trends (Figure 4F). Regarding methylation and transcriptomic values, no IDs were detected in Naxos for both promoter and genebody, whereas only one ID (promoter) and two IDs (genebody) were found at both examined stages in Lakoma (Figures 4G, 4H). One gene ID for Naxos (Soltu.Atl.06_3G007420; Fe superoxide dismutase) and seven for Lakoma (etc. Soltu.Atl.01_2G024990; GDSL-like lipase/acylhydrolase superfamily protein, Soltu.Atl.03_3G018760; peroxidase superfamily protein) showed transcriptomic and proteomic abundance in both stages (Figure 4I).

**Figure 4.**
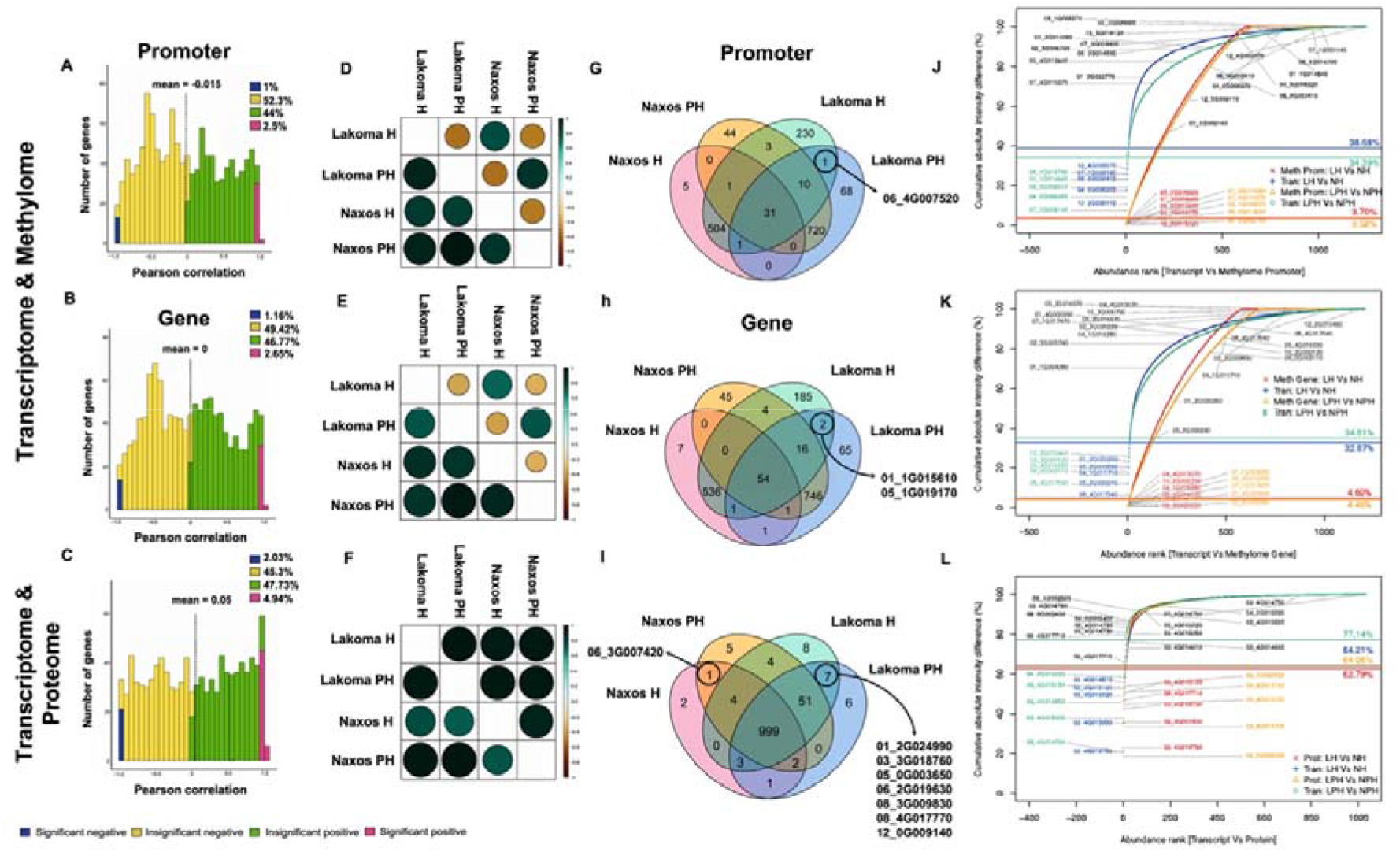
Pairwise transcriptome-based co-expression analysis across methylation, transcriptome and proteome datasets. (A, B, C) Pearson correlation values’ distribution for each omic dataset integration. Methylome-to-transcriptome Pearson correlation heatmaps for (D) promoter (E) gene and (F) transcriptome-to-proteome Pearson correlation heatmap. Venn diagrams of each integrated analysis for transcriptome-to-promoter methylation (G), gene body methylation (H) and proteome (I). Intensity plots displaying significant cumulative difference between Naxos and Lakoma in both stages based on transcriptome-promoter methylome (J), transcriptome-gene methylome (K) and transcriptome-proteome (L) consensus dataset. H: Harvest; PH: Post-harvest. Data obtained from Supplementary Tables 6-9.

We also highlighted the five most significant abundance shifts in our cross-omics datasets (Figures 4J, 4K, 4L). Most methylated IDs of both promoter and gene, were unique for Naxos and Lakoma, respectively (Figure 4J, 4K). Carboxypeptidase A inhibitor domain containing protein (Soltu.Atl.07_1G009140) and PATATIN-like protein (Soltu.Atl.08_0G002410) decreased their transcriptional activity in Naxos during harvest and post-harvest. Accordingly, a zinc finger (C2H2 type) family protein (Soltu.Atl.04_0G006320) gene increased its expression in both stages (Figure 4J). Naxos cultivar showed decreased genebody methylation of serine carboxypeptidase-like (Soltu.Atl.05_2G016570) in harvest, as 4-phosphopantetheine adenylyltransferase (Soltu.Atl.02_3G005740) was decreased in both promoter and genebody at post-harvest (Figures 4J, 4K). It is noticeable that the protein abundance and gene expression of the potato type II proteinase inhibitor family containing domain (Soltu.Atl.03_4G015120) was negatively affected in Naxos at both stages (Figure 4K). Kunitz family trypsin and protease inhibitor protein (Soltu.Atl.03_4G015020), as well as trypsin and protease inhibitor containing domain protein (Soltu.Atl.03_4G014750), were among the top differentiated IDs on a transcript level and decreased their expression in Naxos at both stages (Figure 4K). As for the top differentiated proteins, serine protease inhibitor (Soltu.Atl.09_4G017710) decreased in Naxos for both stages, whereas acyl transferase/acyl hydrolase/lysophospholipase superfamily protein (Soltu.Atl.08_0G002400) and trypsin and protease inhibitor containing domain protein (Soltu.Atl.03_4G014790) increased in Naxos at harvest (Figure 4K).

### Triple multi -omics approaches magnify the possibility of tuber biomarkers detection

Aiming to provide a pipeline integrating data from multiple omics layers, we apply analytical tools, such as correlation downstream analysis and causal models, to identify and characterize the PGI-driver biomarkers. Initially, Pearson correlation was used to examine the linear relationship of the consensus gene IDs within methylome (CHH, promoter or genebody), transcriptome, and proteome (Figure 5A, Tables S10-S11). We then illustrated the changes in tubers of Naxos via heatmap and cluster analysis (Figure 5B), presenting distinct differences among -omics datasets. Analyzing the outcome in detail, we have focused on clusters with a steadily increasing pattern across -omics data in both methylome CHH, promoter or gene body at harvest and post-harvest. For instance, UDP-Glycosyltransferase (Soltu.Atl.05_3G001570), Glycosyl hydrolase (Soltu.Atl.03_2G010490), and zinc finger (C2H2 type) (Soltu.Atl.04_0G006320) increased mainly their levels of proteome and transcriptome in Naxos tubers, with no effect on promoter’s methylome, while this pattern was not observed on gene body methylome CHH-DMR, but in CHG-DMR with 4-alpha-glucanotransferase (Soltu.Atl.02_4G000640). In contrast, lipoxygenase (Soltu.Atl.08_3G003490) has been found to decrease the levels of methylome promoter, proteome, and transcriptome in Naxos tubers. Notably, glutathione *S*-transferase (Soltu.Atl.09_0G002590) has been increased CHH-DMR methylation in both promoter and gene body at harvest followed by a decrease in transcriptome and proteome at both stages (Figure 5B). Following gene ontology (GO) classification the datasets from CHH-DMR methylation in promoter or gene body, transcriptome, and proteome have been enriched to unravel the major groups of genes/proteins that are involved. This approach evidenced that protein- and ATP-binding were the most enriched molecular functions in the triple datasets (Figure 5C).

**Figure 5.**
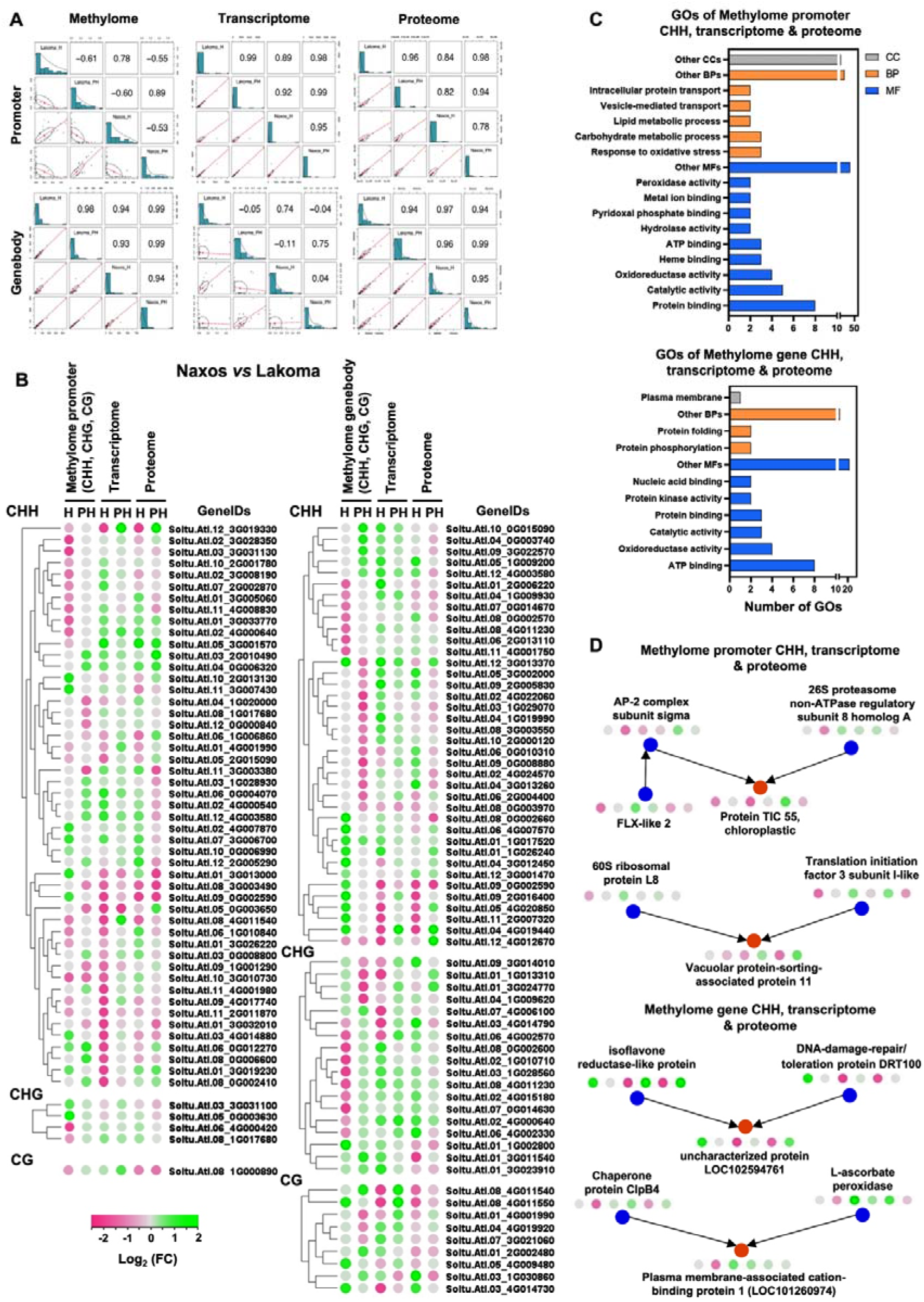
Methylome- (promoter / gene body), transcriptome-, and proteome-based interactions of tubers in the regions of Naxos and Lakoma at harvest (H) and post-harvest (PH). (A) Pearson coefficient was calculated to assess the correlation of the consensus gene IDs in methylome, transcriptome, and proteome only in triplets with values greater than 1 within transcriptomic and proteomic data. (B) Heatmap and clustering of gene IDs from merged datasets of methylome promoter or genebody, transcriptome, and proteome, in the basis of CHH-DMR, CHG-DMR, and CG-DMR. (C) Gene ontology (GO) enrichment analysis of methylome promoter (n=49, CHH) or gene body (n=38, CHH), transcriptome, and proteome. (D) A causal Bayesian network was constructed to detect causality among variables of -omics datasets with consensus gene IDs. Data obtained from Tables S10-S11.

In this study, we employed the dynamics of causal models between variables of consensus genes from the triple datasets to determine possible causal relationships among genes or proteins IDs. Four causal relationships were determined (three of them V-type). Among them, the isoflavone reductase-like protein (Soltu.Atl.04_4G019440) along with the DNA-damage-repair/ toleration protein DRT100 (Soltu.Atl.11_2G007320), as well as the chaperone protein ClpB4 (Soltu.Atl.06_2G004400) with the L-ascorbate peroxidase (Soltu.Atl.09_2G005830), were the cause of the uncharacterized protein LOC102594761 (Soltu.Atl.05_4G020850) and the plasma membrane-associated cation-binding protein 1 (Soltu.Atl.10_2G000120), respectively (Figure 5D).

### Microbials coupled with -omics data as a novel approach to enhance biomarker discovery

As a final step of the present work, the PGI-related molecular changes jointly detected by triple multi -omics analysis (Figures 5, 6) were integrated with metagenomics, enormously expanding the possibility to identify potential biomarkers in PGI foods. To achieve this, a combination of multi-omics datasets (methylome, transcriptome, and proteome) with the metagenome dataset, weighted network analysis (WGCNA) was employed. Using this approach, 30 modules were generated from datasets (Table S12). Initially, we performed a Pearson correlation analysis (Figure 6A) included all pairwise comparisons of the 30 modules corresponding to the five datasets (methylome promoter/gene, transcriptome, proteome, and the microbials; Table S13). This analysis resulted in 232 (53.3% of all comparisons) pairwise comparisons being higher than 0.5 in absolute value (115 positive and 117 negative correlations). Thereafter, a correlation network illustration was constructed (Figure 6B). The edges between nodes (correlations higher than |0.5|), the thickness (absolute value), and the size of the node (degree of centrality) were incorporated in this network. In the next step, the positive correlation of two separate groups that included the microbial modules M1 and M2 was determined and marked the solid lines by red and purple color, respectively (Figure 6B). Then, module eigengenes were used to determine patterns across modules, especially associated with Naxos. Within each group (M1, M9, M15, M22, M24, M28), (M2, M11, M12, M17, M25, M27), an increase of eigengenes in Naxos based on *z*-score at harvest and post-harvest, respectively, was evident (Figure 6C, Table S14).

**Figure 6.**
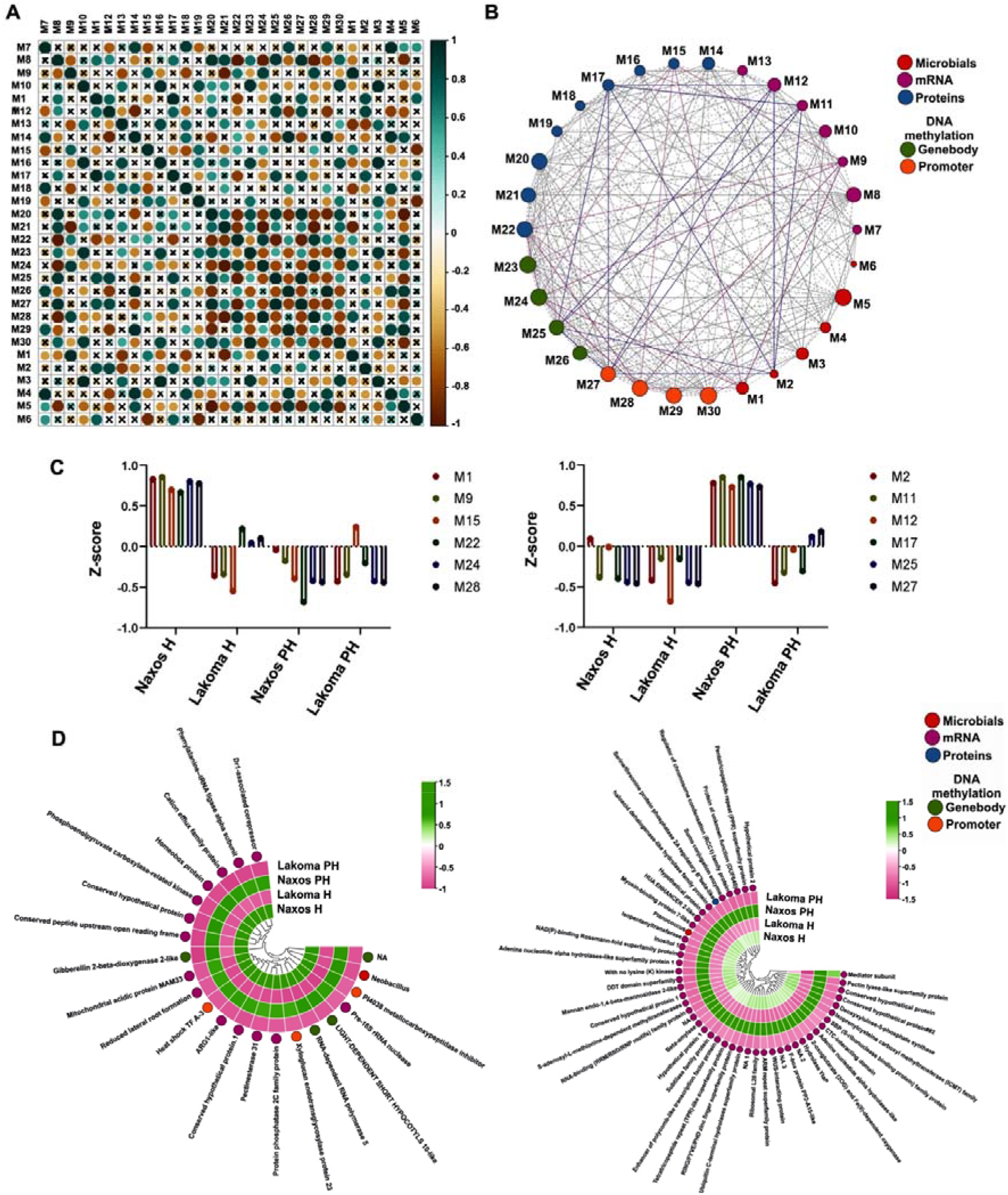
Weighted correlation network analysis (WGCNA) of microbials, mRNAs, proteins, and DNA methylation in Naxos vs Lakoma at harvest (H) and post-harvest (PH). (A) Pearson correlation of 30 modules. The magnitude of the correlation is depicted in both the color and size of the spheres. Correlations which were lower than 0.5 in absolute value are marked with an ‘x’. (B) Network illustration of microbials (Modules M1-M6), mRNAs (Modules M7-M13), proteins (Modules M14-M22), and DNA methylation (genebody modules M23-M26, promoter modules M27-M30) and positive correlation of M1 and M2 with the rest of modules. The modules are represented by the network nodes. The edges connecting the nodes are displayed only when the nodes are correlated with a Pearson coefficient higher than 0.5 in absolute value. Solid lines correspond to positive correlations and dotted lines correspond to negative correlations. The thickness of the lines reflects the magnitude of the correlation (absolute values). The size of the node indicates the degree of centrality (number of edges drawn from the node). (C) The trend of modules-interest M1, M9, M15, M22, M24, M28, and M2, M11, M12, M17, M25, M27, based on their z-scores. (D) Heatmap of positive correlated (P ≤ 0.01) mRNAs (higher tpm than 2 in stages and areas), proteins, and DNA methylation with specific microbials: *Neobacillus* (M1) and *Planococcus* (M2). Data obtained from Tables S12-S14.

To detect pairwise targeted correlations microbial of interest, we focused on the most abundant microbials (genus levels) of the PGI potatoes (from Naxos) compared to the control ones (from Lakoma). These included *Neobacillus* from M1 and *Planococcus* from M2. For both microbial taxa, it should be highlighted that they were found to be dominant and highly abundant solely in Naxos, regardless the individual field, whilst they were present at very low abundance in Lakoma. A Pearson correlation analysis was employed based on these two genera inside each positively correlated group of modules, respectively. Hence, *Neobacillus* has been positively correlated with 14 transcripts (M9) and seven DNA methylation CHH-DMR-base genes correspond to four gene bodies (M24) and three promoters (M28), while *Planococcus* with 45 transcripts (M11 and M12) and one protein (M17).

The large number of transcripts associated with these microbial taxa is a controversial issue, whereas those related to proteins and DNA methylation are considered more trustworthy due to the higher environmental constancy. In this context, *Neobacillus* is related to hypermethylation of gibberellin 2-beta-dioxygenase 2-like (Soltu.Atl.05_2G019380) and xyloglucan endotransglycosylase protein 23 (Soltu.Atl.03_4G007570) in gene body and promoter, respectively. Similarly, *Planococcus* is related to the protein of serine/threonine protein phosphatase 2A regulatory subunit beta-like (Soltu.Atl.04_3G005110).

## DISCUSSION

Undoubtedly, there is an increasing demand by consumers for reliable labeling of food products, as well as for strict controls by retailers, manufacturers, and governments concerning food safety. The development of fast, accurate, and convenient technologies is of utmost importance to assess and trace the authenticated agricultural products from a certain terroir (Wei et al., 2022). The emerging -omics technologies including genomics, epigenomics, transcriptomics and proteomics, and more recent approaches such as metagenomics have been developed with promising applications towards geographical origination (Balkir et al., 2021). Specifically, metagenomics represents a useful diagnostic tool to identify microbial signatures related to natural ecosystems where plants and animals originate from (Iquebal et al., 2022). Pairing the usual -omic tools of Foodomics with microbe-based methods such as amplicon metagenomics, represents a novel and promising technique to improve our understanding on the role of plant-microbe in deciphering plant performance under distinct terroirs (Nerva et al., 2022). Despite the breakthrough of multi-omics studies that brought food research into a new era, the association of Food products with PGI traits has not been conducted yet. Furthermore, to the best of our knowledge, no broad-scale quantitative and integrative analysis of transcriptomes, epigenomes, and proteomes has been performed yet, that would enable the application of the ‘Foodomics’ approach in plant-derived foods. The lack of such studies mostly relies on the difficulties that are raised with the integration of the multiple -omics approaches due to the need of demanding bioinformatics tools and mathematical models required for the accurate quantification and characterization of such large amount of data. In the present study, we achieved the integration of multiple -omics technologies through the application of analytical tools, such as correlation downstream analysis and causal models, aiming to the identification and characterization of putative PGI-driver biomarkers.

The transcript expression and protein abundance of the potato grown at different environments exhibited distinct changes, confirming previous observation that transcriptome and proteome represents a useful diagnostic tool to identify plant performance under distinct terroirs (Braconi et al., 2021; Capozzi and Bordoni, 2012; Wei et al., 2022). For instance, the sucrose synthase protein which catalyzes the conversion of sucrose into glucose and fructose has been found in higher abundance at both stages of Naxos potatoes. This quality trait may be responsible for the characterization of ‘premium quality’ in Naxos potatoes, since an enhanced sucrose synthase activity has been correlated with an increase in starch level and yield in potatoes (Baroja-Fernández et al., 2009). Another interesting finding is that a Kunitz trypsin inhibitor, which probably acts as a regulator of endogenous proteases and assists in defense against pests and pathogens (Bendre et al., 2018), has been revealed to decrease at both transcript and protein levels through the single and pairwise analysis in Naxos potatoes (Fig. 3 and 4). By pairing the transcriptomic and epigenomic or proteomic tools with microbe-based methods, such as amplicon metagenomics, we uncovered that potato tubers cultivated in the semi-arid region of the island of Naxos, which represents a unique Mediterranean agroecosystem, recruit more beneficial microorganisms, possibly to cope with the unfavorable environmental conditions (Leontidou et al., 2020). For example, hypermethylation of xyloglucan endotransglycosylase protein 23 (modify cell wall (Eklöf and Brumer, 2010)) and gibberellin 2-beta-dioxygenase 2-like (catalyzes gibberellin (Santner and Estelle, 2009) that promoting sprouting in potato tubers (Sonnewald and Sonnewald, 2014)) could be driven by *Neobacillus*, which was found to be a dominant and highly abundant species being present only in Naxos tubers (Fig. 6c). These results are in accordance with the notion that soil microbial biogeography is predominantly governed by regional soil properties, unique for each terroir (Fierer and Jackson, 2006; Genitsaris et al., 2020). It is interesting to note that a key microbe like as *Neobacillus* was preserved during post-harvest storage, thus representing an excellent tool towards authenticating these PGI products (Fig. 1 and 6). The rhizosphere of potatoes has been previously found to be a rich source of *Bacillus* strains possessing plant-growth-promoting properties (Calvo et al., 2010), however the exact role of such strains in improving plant performance, quality or shelf-life remains elusive.

Most authentication techniques for food products have focused on species or varietal identification, as well as on the chemical composition of processed foods (Sentandreu and Sentandreu, 2011). Yet, quality traits of plant products can also be determined by cultivation conditions (climate, location, management systems, soil conditions etc.) (Posner et al., 2008). Importantly, cultivation conditions have been shown to induce DNA methylome changes in a wide variety of plants (Lira-Medeiros et al., 2010; Verhoeven et al., 2010). DNA methylome reflects the potato tubers’ perspective of the growing environment, therefore can serve as an appealing diagnostic biomarker (epimarker) tool for geographical origin of otherwise identical crops (López and Wilkinson, 2015). Our findings demonstrated that hypermethylation of glutathione *S*-transferase (enzymes that is induced by stress (Roxas et al., 1997)) at the gene body and promoter leads to a decline of both protein and transcript levels in Naxos tubers (Fig. 5b). Moreover, the causal analysis revealed a V-type connection, where the cause is the (UP) LOC102594761 (Figure 5D) driving us to conclude that this ID is crucial and needs further analysis to understand the proposed connection about hypermethylation of isoflavone reductase (involved in secondary metabolites biosynthesis (Shoji et al., 2002)), DNA-damage-repair/ toleration protein DRT100 (protect DNA under stress (Fujimori et al., 2014)) and uncharacterized protein (UP) LOC102594761 combined with a decrease in transcripts and proteins at harvest and an increase at post-harvest in Naxos (Figure 5D).

The results from this study are of interest also beyond geographical origin studies. For example, putative epimarkers such as the hypermethylation of DETOXIFICATION 18 (Soltu.Atl.10_4G001390) in Naxos, could be used not only to tag cultivation system and geographical region of origin, but also in more nuanced applications to satisfy the ever-increasing demand of the consumers for high quality food products. These could be either in identifying the tissue of origin in plant products (since different plant tissues have diverse methylation profiles) or other factors affecting post-harvest food quality such as storage, transport, and processing conditions (López and Wilkinson, 2015). Collectively, this innovative broad-scale quantitative and integrative work validated the expression of key gene markers by their protein abundance and identified putative epimarkers as well as key microbes to authenticate a popular PGI product such as Naxos potato. At last, but not at least a novel pipeline was developed towards the establishment of breakthrough approaches towards food characterization and authentication.

### Limitations of the study

Our study represents the first multi-omic approach integrating the transcriptome, the proteome, and the epigenome, with the metagenome of a potato of Protected Geographical Indication, to identify terroir-specific “footprints”. However, as the biomarkers identified following causal-model analysis entirely depend on computational modeling, further experimentation is necessary to validate their biological significance and causality, as well as their stability and persistence over years or different geographic regions. On a bioinformatic note, the lack of polyploid specific genome-guided assemblers able to use more than one reference genome, such as in the case of tetraploid potato, may lead to missing alternative homologous sequences, limiting the potential for in-depth downstream transcriptomics analysis. Conclusively therefore, there are still-existing challenges, both experimental and methodological, in capturing dominant terroir-originated marks to diversify and authenticate Protected Geographical Indication agricultural products that are stable across growing seasons and post-harvest storage.

## Materials and methods

### Potato cultivation and experimental sites

Potatoes, cultivar Spunta (Oldenburger, Assem, Holland), were cultivated in two regions of Greece, i.e., Naxos Island, Aegean Sea, Greece, and Lakoma, Chalkidiki, North Greece (Figure 1A, Table S15), following the same experimental and cultivation protocol (composition analysis of potatoes provided in Table S16). The soils in Naxos were characterized as loamy sand or sandy loam (clay content, 13%; sand content 66%), with relatively high organic matter content (2.1%), and pH 7.6 whereas the soils in Lakoma were characterized as clay loam (clay content, 27%; sand content, 45%), with lower organic matter content (1.4%) and pH 7.9. Soil cultivation, fungicide treatments and water application during dry periods, were carried out in accordance with the common potato production schemes in Greece. During crop growth, plants were regularly monitored for the occurrence of stress, pests, and diseases. The harvest of the tubers was performed early in June 2021 for both collection sites, after foliage desiccation. All tubers were placed in sterile bags at 10 °C and carried to the lab, within 12 hours. Samples for subsequent analyses were snap-frozen in liquid nitrogen and stored at −8□°C. At harvest, there were eight and six samples (pooled tubers from an individual plant, with three biological replicates for each sample), corresponding to different collection sites of Naxos and Lakoma, respectively. The exact sampling locations are provided in Table S15. For the post-harvest experiment, tubers were stored at 10 °C for one-month prior subsequent analyses.

### Transcriptome and whole-genome bisulfite sequencing

#### Library construction

Total RNA of pooled tubers from the eight collection sites of Naxos, and the six collection sites of Lakoma (Table S15), with three biological replicates each, was isolated using TRIzol™ reagent (Invitrogen, CA, USA), followed by rRNA depletion and DNaseI treatment (Qiagen, Hilden, Germany). For each RNA sample at harvest and at post-harvest, a paired-end strand-specific Tru-seq compatible library was constructed following manufacturer’s instructions.

High quality genomic DNA was isolated from the same samples as previously described for the RNASeq experiment, using the CTAB method (Doyle J. J. and Doyle J. L., 1990) to perform whole-genome bisulfite sequencing (WGBS) at Beijing Novogene Technology Co., Ltd. with target sequencing depth at 30×. The isolated DNA was fragmented by sonication to 200-300 bp using a Covaris S220 (Covaris, Woburn, MA, USA), followed by end repair and A-ligation. After ligation to cytosine-methylated barcodes, the DNA fragments were treated twice with bisulfite using an EZ DNA Methylation-Gold™ Kit (Zymo Research, Orange, CA, USA). The libraries were then prepared according to the Illumina standard DNA methylation analysis protocol.

RNA-seq and WGBS libraries were sequenced (Paired-End, 150bp) on the Illumina Novaseq 6000 platform (Illumina, CA, USA) (Novogene, Beijing, China).

#### RNA-seq data analysis

All data generated were aligned to the tetraploid *Solanum tuberosum* reference genome (Atlantic v2.0) via Hisat2 applying the default parameters. Quantification of raw read counts/gene was conducted with HTSeq v0.11.1 (http://www-huber.embl.de/users/anders/HTSeq/), selecting the ‘-s reverse -type=gene’ option. Transcripts Per Million (TPM) was used for the normalization of raw reads. Data were then transformed to log2 and scaled (z-score: mean center divided by standard deviation). Principal Component Analysis was performed for each sample and treatment) using the normalized data with the function ‘prcomp’ in R (version 4.1.0).

#### Whole-genome bisulfte sequencing (WGBS) data analysis

FastQC (fastqc_v0.11.5) was used to perform basic statistics on the quality of raw reads. These sequences, produced by the Illumina pipeline in FASTQ format, were pre-processed through Trimmomatic (Trimmomatic-0.36) software using the parameter (SLIDINGWINDOW: 4:15; LEADING:3; TRAILING:3; ILLUMINACLIP: adapter.fa: 2: 30: 10; MINLEN:36). The low quality (< Q30) data was filtered out, and the filtered high quality sequencing data was mapped to the tetraploid *Solanum tuberosum* reference genome (Atlantic v2.0) by Bismark v0.19.0 (Krueger and Andrews, 2011).

The Bioconductor package DSS (Dispersion Shrinkage for Sequencing) was used to identify differentially methylated regions (DMRs) following default parameter settings with a reduced smoothing size (smoothing span□=□200). According to the distribution of DMRs throughout the genome, genes related to DMRs were defined as DMR-associated genes whose gene body region (from TSS to TES) or promoter region (upstream 2□kb from the TSS) overlapped with DMRs.

#### Quantitative real time (qRT) PCR assay

Total RNA isolated from aliquots of the sequenced samples was reverse transcribed to cDNA using SuperScript™ First-Strand Synthesis System (Invitrogen™ Thermo Fisher Scientific, Inc.). Gene expression profiles of ten genes (Table S17), randomly picked from the dataset, were analyzed by Quantitative real time PCR (qRT PCR) using Luna® Universal qPCR Master Mix (New England BioLabs) in a QuantStudio® 5 Real-Time PCR System (Thermo Fisher Scientific) according to (Xanthopoulou et al., 2021). For gene expression normalization, *EF-1a* was used as reference gene (Tang et al., 2017).

### Proteomics

#### Bottom-up proteomic sample preparation

The protein extracts, obtained from three representative collection sites in Naxos and three in Lakoma, with three biological replicates each one at harvest and at post-harvest, were processed according to the sensitive Sp3 protocol. The cysteine residues were reduced in 100 mM DTT and alkylated in 200 mM iodoacetamide (Acros Organics). 20 ug of beads (1:1 mixture of hydrophilic and hydrophobic SeraMag carboxylate-modified beads, GE Life Sciences) were added to each sample in 50% ethanol. Protein clean-up was performed on a magnetic rack. The beads were washed twice with 80% ethanol and once with 100% acetonitrile (Fisher Chemical). The captured-on beads proteins were digested overnight at 37°C under vigorous shaking (1200 rpm, Eppendorf Thermomixer) with 1 ug Trypsin/LysC (MS grade, Promega) prepared in 25 mM Ammonium bicarbonate. The next day, the supernatants were collected, and the peptides were purified using a modified Sp3 clean up protocol and finally solubilized in the mobile phase A (0.1% Formic acid in water), sonicated and the peptide concentration was determined through absorbance at 280nm measurement using a nanodrop instrument.

#### LC-MS/MS Analysis

Samples were analyzed on a liquid chromatography tandem mass spectrometry (LC-MS/MS) setup consisting of a Dionex Ultimate 3000 nanoRSLC coupled inline with a Thermo Q Exactive HF-X Orbitrap mass spectrometer. Peptidic samples were directly injected and separated on a 25 cm-long analytical C18 column (PepSep, 1.9μm3 beads, 75 μm ID) using an one-hour long run, starting with a gradient of 7% Buffer B (0.1% Formic acid in 80% Acetonitrile) to 35% for 40 min and followed by an increase to 45% in 5 min and a second increase to 99% in 0.5min and then kept constant for equilibration for 14.5min. A full MS was acquired in profile mode using a Q Exactive HF-X Hybrid Quadrupole-Orbitrap mass spectrometer, operating in the scan range of 375-1400 m/z using 120K resolving power with an AGC of 3x 106 and maximum IT of 60ms followed by data independent acquisition method using 8 Th windows (a total of 39 loop counts) each with 15K resolving power with an AGC of 3x 105 and max IT of 22ms and normalized collision energy (NCE) of 26.

#### Proteomic Data Analysis

Orbitrap raw data from the 35 protein samples (one has failed) were analyzed in DIA-NN 1.8 (Data-Independent Acquisition by Neural Networks) through searching against the Atlantic v2.0 (http://spuddb.uga.edu/phased_tetraploid_potato_download.shtml) using the library free mode of the software, allowing up to two tryptic missed cleavages and a maximum of three variable modifications/peptide. A spectral library was created from the DIA runs and used to reanalyze them (double search mode). DIA-NN search was used with oxidation of methionine residues and acetylation of the protein N-termini set as variable modifications and carbamidomethylation of cysteine residues as fixed modification. N-terminal methionine excision was also enabled. The match between runs feature was used for all analyses and the output (precursor) was filtered at 0.01 FDR and finally the protein inference was performed on the level of genes using only proteotypic peptides. The generated results were processed statistically and visualized in the Perseus software (1.6.15.0).

### Metagenomics

#### DNA Extraction, Amplification, and Sequencing

Nearly 200 grams of 72 samples obtained from potato tuber-sphere at harvest or after one-month post-harvest storage, from all the individual collection sites (Table S15), was used for the microbial mapping of the two different regions. High quality DNA was isolated with the DNeasy PowerSoil Pro Kit (QIAGEN, Carlsbad, USA), following the manufacturer’s instructions and stored at −80°C. Amplification of the 16S rRNA gene was performed using an Applied Biosystems® QuantStudio® 5 Real-Time PCR System (Thermo Fischer Scientific, Waltham, MA, USA), using a LongAmp Hot Start Taq 2x Master Mix (M0533S, New England Biolabs), and 16S barcoded primers.

The 16S Barcoding Kit 1-24 (SQK-16S024, Oxford Nanopore Technologies, UK) was used for sequencing the 16S ribosomal gene and creating the libraries. PCR products were purified with Agecount AMPure XP beads (Beckman Coulter, USA), whilst the quantification was performed using Qubit 4 Fluorometer and the dsDNA HS Assay Kit (Thermo Fisher Scientific, USA). The 72 libraries were created in accordance with the manufacturer’s instructions and loaded on a MinION R9.4.1 flow cell (FLO-MIN106) on the MinION Mk1C (Oxford Nanopore Technologies, UK). For data acquisition, MINKNOW software ver. 1.11.5 (Oxford Nanopore Technologies) was employed.

#### Sequencing Data Processing and Analysis

MinION™ sequence reads (i.e., FAST5 data) were converted into FASTQ files by using Guppy software (version 5.0.17) (Oxford Nanopore Technologies). To remove reads derived from humans, EPI2ME 16S pipeline software was used. The unmatched reads to the human genome were considered as reads obtained from bacteria.

Bacterial communities were identified through the same software (EPI2ME), which is based on Nextflow (di Tommaso et al., 2017), that enables scalable and flexible scientific analysis (Delegou et al., 2022). In order to classify the DNA sequences from microbial samples, the Centrifuge software was used (Kim et al., 2016), which is based on the Burrows-Wheeler Transformation (BWT) and the Ferragina-Manzini (FM) index, that enables timely and precise metataxonomic analysis. Operational taxonomic units (OTU tables) by matching the NCBI taxa IDs to lineages and counting the number of reads per NCBI taxa ID.

Alpha – diversity was calculated, using the “vegan” package, while beta – diversity was assessed, applying the “vegan” and “betapart” packages (Baselga and Orme, 2012), all in R studio software. Principal component analysis (PCA) and Non-metric Multi-dimensional Scaling (NMDS) were conducted via “vegan” (https://github.com/vegandevs/vegan) and “graphics” (https://rdrr.io/r/graphics/graphics-package.html) packages. In addition, analysis of similarities (ANOSIM) was also performed, using the vegan package. In order to run the hierarchical clustering algorithm, multiple dendrograms by chaining were performed, using the “tidyverse” (https://www.tidyverse.org/) and “dendextend” packages (Galili, 2015). A heatmap based on the relative abundance of OTUs was generated using the “gplots” package (https://cran.r-project.org/web/packages/gplots/index.html), while stacked bar charts were performed, integrating the top 10 most abundant genera and species between tuber samples at harvest and at post-harvest. Finally, the linear discriminant analysis (LDA) effect size (LEfSe) analysis was conducted (Segata et al., 2011) via Galaxy software (https://huttenhower.sph.harvard.edu/galaxy/), in an effort to characterize the microbial variance between the unique categories and determine possible biomarkers for each one.

### Multi-omics analysis

#### Dual approach

The analysis was separately performed for the pairs corresponding to the same gene IDs in transcriptome and methylome promoter or genebody, and transcriptome/proteome. Only pairs which exhibited valid values for all tissues at both levels were considered. Of these, for the transcript/protein, only pairs with values greater than 1 in at least one out of the four groups at both transcriptomic and protein levels were further assessed (the methylome values were between 0 and 1), resulting in 1247 transcript/methylome promoter pairs, 1206 transcript/methylome gene pairs, and 1033 transcript/protein pairs. The Pearson coefficient was used to assess the correlation in all three dual comparisons, across and between groups, respectively. The ranking of the absolute mean intensity differences in pairwise comparisons (Naxos *vs* Lakoma (Harvest), Naxos *vs* Lakoma (Post-Harvest)) was used as well.

#### Triple approach

The Pearson coefficient was further employed to assess the correlation between tissues for the triplets corresponding to consensus gene IDs in methylome promoter/transcriptome/proteome (n=49), and methylome genebody/transcriptome/proteome (n=38). Only triplets with valid values for the stages and areas were considered, which exhibited values greater than 1 in at least one out of the stages or areas at both transcriptomic and protein levels. Next, the focus was in identifying causal relations between methylome/transcriptome/proteome triplets. To this end, the constrained-based PC algorithm, was employed (“pcalg” R package, https://cran.r-project.org/web/packages/pcalg/index.html), which is used to estimate the causal structure induced by a causal Bayesian network. For each pair of variables (X, Y) in a dataset, the PC algorithm evaluates their independence, conditioning on all subsets of all the remaining variables. If their association is persistent, it is considered to be causal. The output is a network represented by a Markov equivalence class of the Directed Acyclic Graph (DAG), with a structure consistent with the results of the tests of independence. It is assumed that causal sufficiency holds2, which implies that for every pair of measured variables, all their consensus direct causes are also measured. A directed edge between X and Y exists, if and only if, the variables are conditionally dependent given S, for all possible subsets S of the remaining nodes. In particular, the “pc” R function was used to estimate the equivalence class of the DAG, under the Markov assumption that the distribution of the observed variables is faithful to a DAG3. All genes exhibited continuous values, thus, the function “gaussCItest” was employed to perform the conditional independence tests.

#### Dataset approach

For each omics dataset (and microbials), weighted gene co-expression network analysis (WGCNA) was employed (“WGCNA” package in R, https://horvath.genetics.ucla.edu/html/CoexpressionNetwork/Rpackages/WGCNA/), to identify data clusters (modules) across areas and stages. The “blockwiseConsensusModules” function was used with minimum module size=30, module detection sensitivity=2, and cut height for module merging=0.25). Next, the eigengenes of the modules were used to assess the correlation among all modules. Module eigengenes are the module representatives and defined as the first principal component of the expression matrix for each module. A module eigengene correlation network was developed as well, with nodes representing the modules, and edges representing all the correlations between the nodes with absolute value higher than 0.5. All the analyses were performed with R Version 4.1.0.

## Data availability

Raw data of RNASeq, Bisulfite-Seq and Metagenome were deposited in the National Centre for Biotechnology Information (NCBI) Sequence Read Archive (SRA) under BioProject accession numbers: PRJNA855343, PRJNA855343 and PRJNA854325, respectively. The mass spectrometry proteomics data have been deposited to the ProteomeXchange Consortium via the PRIDE partner repository (Perez-Riverol et al., 2022) with the dataset identifier PXD035074.

## Acknowledgements

This research was funded by European Regional Development Fund (ERDF), through the Operational Program “Southern Aegean” 2014–2020, entitled “Enhancement of quality and nutritional traits of Naxos potatoes using omics-technologies. Acronym: GrEaTest-Potatoes” (0040991). The implementation of the doctoral thesis was co-financed by Greece and the European Union (European Social Fund-ESF) through the Operational Programme «Human Resources Development, Education and Lifelong Learning» in the context of the Act “Enhancing Human Resources Research Potential by undertaking a Doctoral Research” Sub-action 2: IKY Scholarship Programme for PhD candidates in the Greek Universities. We acknowledge support of this work by the project “The Greek Research Infrastructure for Personalised Medicine (pMedGR)” (MIS 5002802) which is implemented under the Action “Reinforcement of the Research and Innovation Infrastructure”, funded by the Operational Programme ‘Competitiveness, Entrepreneurship and Innovation’; (NSRF 2014-2020) and co-financed by Greece and the European Union (European Regional Development Fund).

## Authors contribution

I.G. and I.M. conceived and designed the experiment; I.G and I.M planned the structure of the paper; AB and MM prepared the first draft manuscript; A.B., C.B., M.S., G.S, T.M. and M.G. analyzed the data; MG carried out the causal model analysis; A.B., M.M., A.D., M.S., G.T., C.S., A.X., G.S., I.G., L.A., A.M., I.N-O., G.T., I.F. and C.B. contributed to data acquiring. All authors contributed to consensus results interpretation and revised the final manuscript.

## Supplemental information

Table S1. Number of reads and operational taxonomic units (OTUs) for bacterial communities of *S. tuberosum* at harvest and post-harvest.

Table S2. Summary of whole genome DNA bisulfite sequencing data at harvest and post-harvest.

Table S3. Statistics of raw, clean and mapped reads from RNA sequencing at harvest and post-harvest.

Table S4. Overview of RNA sequencing data at harvest and post-harvest.

Table S5. Differential expression analysis of proteomics data at harvest and post-harvest.

Table S6. Integration of transcriptomic (gene) and methylation data to perform dual co-expression analyses. at harvest and post-harvest.

Table S7. Integration of transcriptomic (promoter) and methylation data to perform dual co-expression analyses. at harvest and post-harvest.

Table S8. Integration of transcriptomic and proteomic data to perform dual co-expression analyses. at harvest and post-harvest.

Table S9. Combination of multi-omics dataset at harvest and post-harvest.

Table S10. Methylome genebody, transcriptome and proteome interaction at harvest and post-harvest.

Table S11. Methylome promoter, transcriptome and proteome interaction at harvest and post-harvest.

Table S12. Modules generated from weighted network analysis (WGCNA) at harvest and post-harvest.

Table S13. Pearson correlation between targeted microbial genus and -omics data at harvest and post-harvest.

Table S14. Module eigenvalues at harvest and post-harvest.

Table S15. Coordinates for the sampling locations, as well as the samples used for each - omic experiment.

Table S16. Composition analysis of *S. tuberosum*.

Table S17. Gene expression profiles of ten genes analyzed by Quantitative real time (qRT).

